# GEMME: a simple and fast global epistatic model predicting mutational effects

**DOI:** 10.1101/543587

**Authors:** Elodie Laine, Yasaman Karami, Alessandra Carbone

## Abstract

The systematic and accurate description of protein mutational landscapes is a question of utmost importance in biology, bioengineering and medicine. Recent progress has been achieved by leveraging on the increasing wealth of genomic data and by modeling inter-site dependencies within biological sequences. However, state-of-the-art methods require numerous highly variable sequences and remain time consuming. Here, we present GEMME (www.lcqb.upmc.fr/GEMME), a method that overcomes these limitations by explicitly modeling the evolutionary history of natural sequences. This allows accounting for all positions in a sequence when estimating the effect of a given mutation. Assessed against 41 experimental high-throughput mutational scans, GEMME overall performs similarly or better than existing methods and runs faster by several orders of magnitude. It greatly improves predictions for viral sequences and, more generally, for very conserved families. It uses only a few biologically meaningful and interpretable parameters, while existing methods work with hundreds of thousands of parameters.

## Introduction

Understanding which and how genetic variations affect proteins and their biological functions is a central question for bioengineering, medicine and fundamental biology. In these fields, a fast and accurate assessment of the effects of every possible substitution at every position in a protein sequence (full single-site mutational landscape) or of combinations of mutations (pairs, triplets…) would allow to reach some level of control over proteins, needed to improve the treatment of diseases, the design of new proteins and the synthesis of molecular libraries. Deep mutational scans [1] or multiplexed assays for variant effects (MAVEs) [2] have enabled the full description of the mutational landscapes of a few tens of proteins (see [3] for a list of proteins and associated experiments). They have revealed that a protein contains a relatively small number of positions highly sensitive to mutations, where almost any substitution induces highly deleterious effects [4, 5]. Although these methods represent major biotechnological advances, they remain resource intensive and are limited in their scalability. Moreover, the measured phenotype and the way it is measured vary substantially from one experiment to another, making it difficult to compare different measurements and/or proteins [6]. These limitations call for the development of efficient and accurate computational methods for high-throughput mutational scans.

Many computational methods predicting mutational effects exploit information coming from protein sequences observed in nature [3, 7, 8, 9, 10, 11, 12, 13, 14, 15]. They rely on the assumption that rarely occurring mutations induce deleterious effects. Most of them start from a multiple sequence alignment and treat each position in the alignment independently from the others to compute frequencies of occurrence. However, the amino acid residues comprising a protein are inter-dependent, and the effect of a mutation depends on the amino acids present at other positions, a phenomenon referred to as ‘epistasis’ [16, 17]. By leveraging on the increasing wealth of genomic data, recent developments have enabled modeling inter-dependencies between positions and have significantly improved the accuracy of mutational effects predictions [3, 7, 8, 9]. Specifically, some statistical methods estimate couplings between pairs of positions [7, 8]. They are very accurate in identifying a few strong direct couplings responsible for the whole co-variability observed in homologous sequences and corresponding to physical contacts in protein structures [18, 19]. In the context of mutational outcome prediction, however, the usage made of pairwise couplings is fairly different [7, 8]. It is the ensemble of all couplings, between a given position and all other positions, that is relevant, not the identification of a few high ones. Hence, pairwise couplings are used as a proxy to capture the influence of the whole sequence context on a particular position. Nevertheless, the information remains local and the explicit calculation of higher-order couplings is computationally intractable. To circumvent this issue, a deep latent-variable model was proposed where the global sequence context is implicitly accounted for by coupling the observed positions to latent (‘hidden’) variables [3]. The model is fully trained on each studied protein family to generate sequences likely to belong to the family. Deviations between outputs and inputs are then used as estimates of the mutational effects. Biologically meaningful constraints are included in the model to ease interpretability and improve predictions. Although the model sometimes achieves very high correlation with experimental measurements, its performance is highly disparate from one protein family to another. Both pairwise couplings based and deep latent-variable based methods require a large diversity in the available sequence data and are computationally costly.

In this work, we present a fast, scalable and simple algorithm to predict mutational effects by explicitly modeling inter-dependencies between all positions in a sequence, thereby accounting for global epistasis. Key to our approach is the notion of an evolutionary history relating the sequences observed today in nature. We view each sequence as an evolutionary solution selected with respect to a mutation of interest. Our algorithm infers the evolutionary relationships relating natural sequences by quantifying their global similarities. These relationships, encoded in a tree, are used to determine the extent to which each position is *conserved* in evolution, and to estimate the *evolutionary fit* required to accommodate mutations. Our notion of conservation is inspired from that of evolutionary trace [20, 21]. More precisely, we estimate the degree of conservation of a position by computing the evolutionary age of the amino acid at that position over many trees relating homologous sequences. Since the trees represent the divergence of the entire sequences along evolution, the conservation degree of each position is influenced by all other positions. Consequently, this measure goes beyond statistical measures of site-independent conservation and measures of coevolution extracting signals from pairs of positions. It is computed by the Joint Evolutionary Trees (JET) method [22], and it proved to be useful in the identification of protein interfaces [22, 23, 24]. We use the computed conservation degrees to weight positions, the rationale being that changes occurring at more conserved positions likely have bigger impacts on the protein’s function [25]. Then, to be able to discriminate between different substitutions occurring at a given position, we combine two quantities. The first one is the relative frequency of occurrence of the mutation, relying on physico-chemical similarities rather than amino acid identities. The second one is the minimum evolutionary fit required to accommodate the mutation. Namely, we estimate how far one has to go in the evolutionary tree to observe a natural sequence displaying the mutation. We show that our algorithm overall performs on-par with the nonlinear latent-variable model of DeepSequence [3] and better than the pairwise epistatic model of EVmutation [7]. These are currently the best performing methods of the state of the art [3]. Importantly, our approach achieves greatly improved predictions for very conserved protein families. This property makes it the only method suitable to treat viral sequences. Beyond prediction assessment, we provide a clear readout of the contribution of epistasis by looking at how sequences observed in nature are spread and evolutionary related. Our method is implemented as a fully automated tool, Global Epistatic Model for predicting Mutational Effects (GEMME), available as a downloadable package and as a webserver at: www.lcqb.upmc.fr/GEMME/. We show that GEMME is faster than DeepSequence and EVmutation by several orders of magnitudes. It makes possible the systematic study of pairs or triplets of mutations appearing sequentially in time and associated to drug resistance, especially in viruses. It could help in taking informed decisions regarding patients treatment and public health by enabling real-time analysis of pathogenic sequence data [26].

Our results emphasize the usefulness of the information encoded in the way certain positions, because of their functional importance, are segregated along the topology of evolutionary trees. A functionally important position is expected to be associated with one or several subtrees of ancient origin and homogeneous with respect to that position (all sequences in the subtree display the same amino acid). This type of patterns is essentially what our measure of conservation captures. We demonstrate that this notion of conservation, combined with site-independent frequencies, is sufficient to accurately predict mutational effects, and that the notion of coevolution in not required to address this problem. Hence, this work paves the way to new ideas and developments in sequence-and evolutionary-based mutational outcome prediction.

## Results

### Evolutionary-informed model globally accounts for epistasis

Sequences observed in nature have been selected for function through evolution. Hence, they can inform us on the constraints underlying evolutionary processes and help us estimate mutational effects. To assess the impact of a given mutation at a given position in a query sequence, we look at an ensemble of sequences homologous to that query (**Figure 1a**). These sequences can be organized in a tree based on their global similarities (**Figure 1b**). The topology of the tree reflects the evolutionary relationships between the sequences. Our main contribution is to extract conservation patterns in line with the topology of the tree and use them to determine the extent to which a mutation will be deleterious for the function of the query protein. For this, the first step of our approach consists in estimating the biological importance of each residue in the query by computing its evolutionary conservation (**Figure 1c**, first color strip). Highly conserved positions are likely important for the protein stability and/or function and thus likely sensitive to changes. For each position in the query, we look at the level in the tree where the amino acid at that position appeared and remained conserved thereafter. To illustrate this notion of conservation, we consider a toy example with two positions, namely *i* and *j* displaying *S* and *G*, respectively (**Figure 1a-b**). In the tree, one can observe that the amino acid *G* at position *j* was fixed much earlier in evolution compared to *S* at position *i* (**Figure 1b**, compare grey rectangles between the two panels). Hence, we will assign a much higher conservation degree to the former than to the latter. Since the tree is inferred from global similarities between entire sequences, the conservation degree of a given position accounts for the way all other positions have diverged along evolution. Here, we deal with a potentially large number of sequences, and the reconstruction of a unique tree relating all of them may lead to an unreliable topology. To cope with this issue, we sample the initial ensemble of sequences to extract representative subsets and reconstruct trees starting from these subsets (see *Methods*). Conservation degrees are averaged over all reconstructed trees to get statistically significant estimates. We then use them to compare mutations occurring at the same position, and to compare different positions.

**Figure 1:**
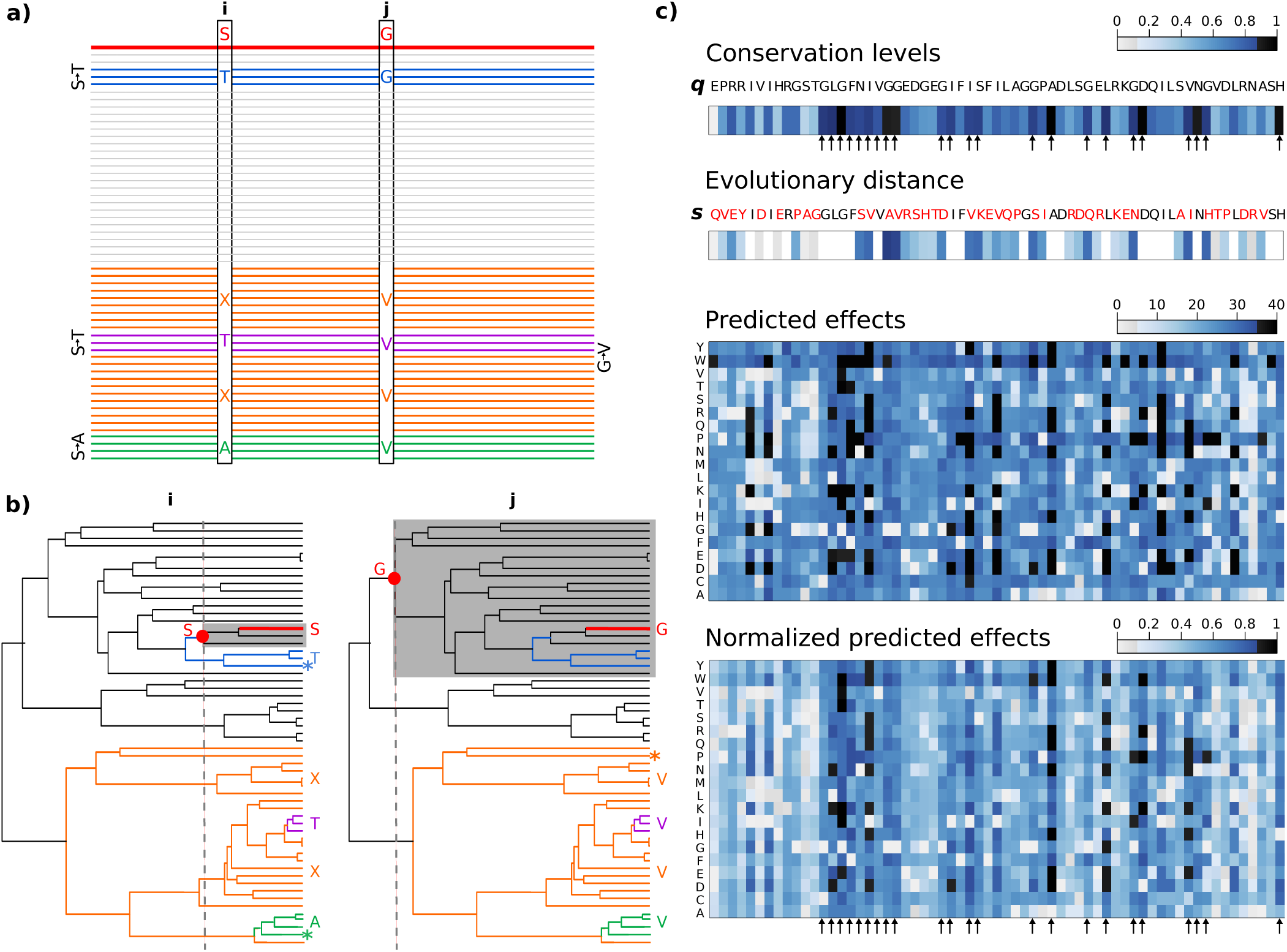
Principle of the method. **(a)** Ensemble of sequences related to a query sequence, on top and in red. The query displays a serine (S) at position *i* and a glycine (G) at position *j*. Some sequences are colored according to the amino acids they display at the two positions: T-G in blue, T-V in purple, X-V in orange (X stands for any amino acid, except for T and A) and A-V in green. **(b)** Tree representing the evolutionary relationships between the related sequences. The color code is the same as in (a). Information concerning positions *i* and *j* is reported on the left and on the right, respectively. The red dots and dotted grey lines indicate the levels where S and G appeared at positions *i* and *j* and remained conserved thereafter. The associated subtrees are highlighted by grey rectangles. The stars indicate the closest sequences to the query displaying the S-to-T mutation at *i* (left, in blue), the S-to-A mutation at *i* (left, in green) and the G-to-V mutation at *j* (right, in orange). **(c)** Workflow of the method applied on the third PDZ domain of PSD95 (DLG4). The color strip on top gives conservation levels computed for the query sequence *q*. Positions highlighted by arrows are highly conserved. A homologous sequence *s* is displayed below, with its mutations highlighted in red. The second color strip indicates the squared conservation levels for the positions of the mutations. The two matrices give the predicted effects and normalized predicted effects, respectively, for all possible substitutions at all positions in *q*.

To compare different mutations at a given position, we introduce the notion of evolutionary fit. It reflects the amount of changes required to accommodate a mutation over the entire sequence. We estimate it by looking at how far natural sequences displaying the mutation are from the query sequence in the evolutionary tree. Our working hypothesis is that the more distant these sequences, the more deleterious the mutation. For the sake of simplicity, let us consider two mutations at position *i*, namely S-to-T and S-to-A, which we want to compare, and see how they are associated to changes at another position *j* (**Figure 1a**). While the S-to-T mutation is sometimes associated to the wild-type G at *j* (sequences in blue), the S-to-A mutation is systematically accompanied by a mutation at *j* (namely G-to-V, sequences in green). Since position *j* is much more conserved than *i*, this will result in sequences bearing S-to-A at *i* being much further away in the tree, with respect to the query, than sequences bearing S-to-T at *i* (**Figure 1b**, compare the locations of the green and blue sequences). Intuitively, this observation suggests that it will be more difficult for the query to accommodate A compared to T at position *i*, and hence that S-to-A will be more deleterious than S-to-T. We can easily generalize this reasoning over two positions to the whole sequence. For this, we define an evolutionary distance between the query *q* and some sequence *s* which explicitly accounts for the conservation degrees of all variable positions between *q* and *s* (see *Methods* and **Figure 1c**). For each studied mutation, we look for the closest sequence to *q* displaying that mutation, and we use its evolutionary distance to estimate the minimal evolutionary fit associated to the mutation. We combine evolutionary fits with site-independent frequencies calculated using a reduced amino acid alphabet to get more precise estimates (see *Methods*).

Then, to be able to compare mutations occurring at different positions, we rely on the hypothesis that more conserved positions will be more sensitive to any mutation than less conserved positions. To implement this idea, we re-weight the predicted mutational effects by the evolutionary conservation degrees (**Figure 1c**, compare the two matrices). As a result, highly deleterious mutations will be mainly found at highly conserved positions (**Figure 1c**, second matrix, dark squares are mainly localized at conserved positions, highlighted by arrows). In our toy example, although the evolutionary distance computed for the G-to-V mutation at *j* is lower than that computed for the S-to-A mutation at *i* (**Figure 1b**, compare the locations of the starred orange sequence on the right and the starred green sequence on the left), the former will be predicted as more deleterious than the latter because position *j* is much more conserved than position *i*.

GEMME’s predictive model globally accounts for epistasis by explicitly looking at the whole sequence context when assessing the effect of a particular mutation. It is applicable to single site mutations and also to combinations of mutations (see *Methods*).

### GEMME predicts mutational outcomes better than state-of-the-art methods, especially for viral sequences

We assessed GEMME’s predictive power against experimental measures collected from 41 high-throughput mutational scans of 33 proteins and 1 protein complex, representing 657 840 mutations (**Supplementary Table S1**). Among them, 38 scans deal with single-site mutations, 1 with pairs of mutations and 2 with multiple mutations. The average Spearman rank correlation between predicted and experimental values is 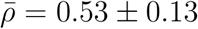. The best agreement is obtained for the bacterial *β*-lactamase, with a correlation of 0.74 (**Fig. 2a**). Compared with the state-of-the-art methods DeepSequence and EVmutation, GEMME performs equally well or better (**Fig. 2a**). Namely, its overall performance are similar to those of DeepSequence [3] (Δ*ρ*_*GEMME-DEEP*_ ≥ 0 in 19/41 scans, with an average of 0.02 ± 0.12 and a median of *-*0.01) and significantly better than those of EVmutation [7] (Δ*ρ*_*GEMME-EV* *mutation*_ ≥ 0 in 32/41 scans, with an average of 0.03 ± 0.05 and a median of 0.03). Importantly, GEMME achieves much higher correlations for the 5 viral sequences of the dataset (**Fig. 2a**, on the right, and **Fig. 2b**, orange dots), up to an impressive Δ*ρ* = 0.5 compared to DeepSequence and Δ*ρ* = 0.1 compared to EVmutation. The input alignments for these proteins display a very low degree of diversity, with more than 60% of sequences sharing more than 60% of identity with the query sequence (**Fig. 3**, underlined proteins). More generally, the lower the diversity of the input alignment, the higher the improvement of GEMME over the two other methods (**Fig. 3**). Consequently, GEMME presents a clear advantage over DeepSequence and EVmutation when the diversity of the available sequence data is low or very low.

**Figure 2:**
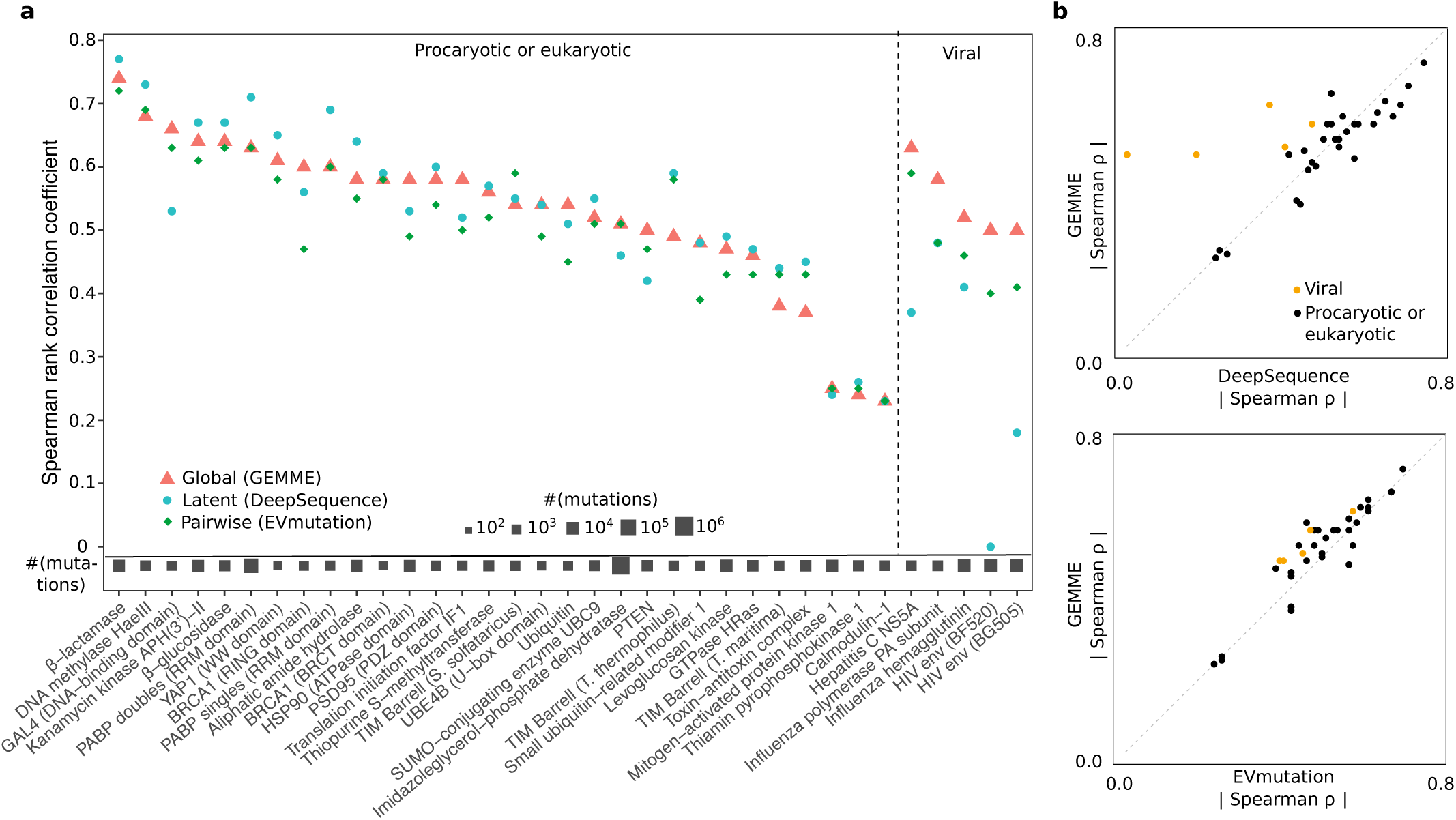
Comparison of predictive performances between GEMME, DeepSequence and EVmutation. Spearman rank correlation coefficients *ρ* between predicted and experimental measures are reported for 35 experiments corresponding to 34 proteins (see *Methods*).

**Figure 3:**
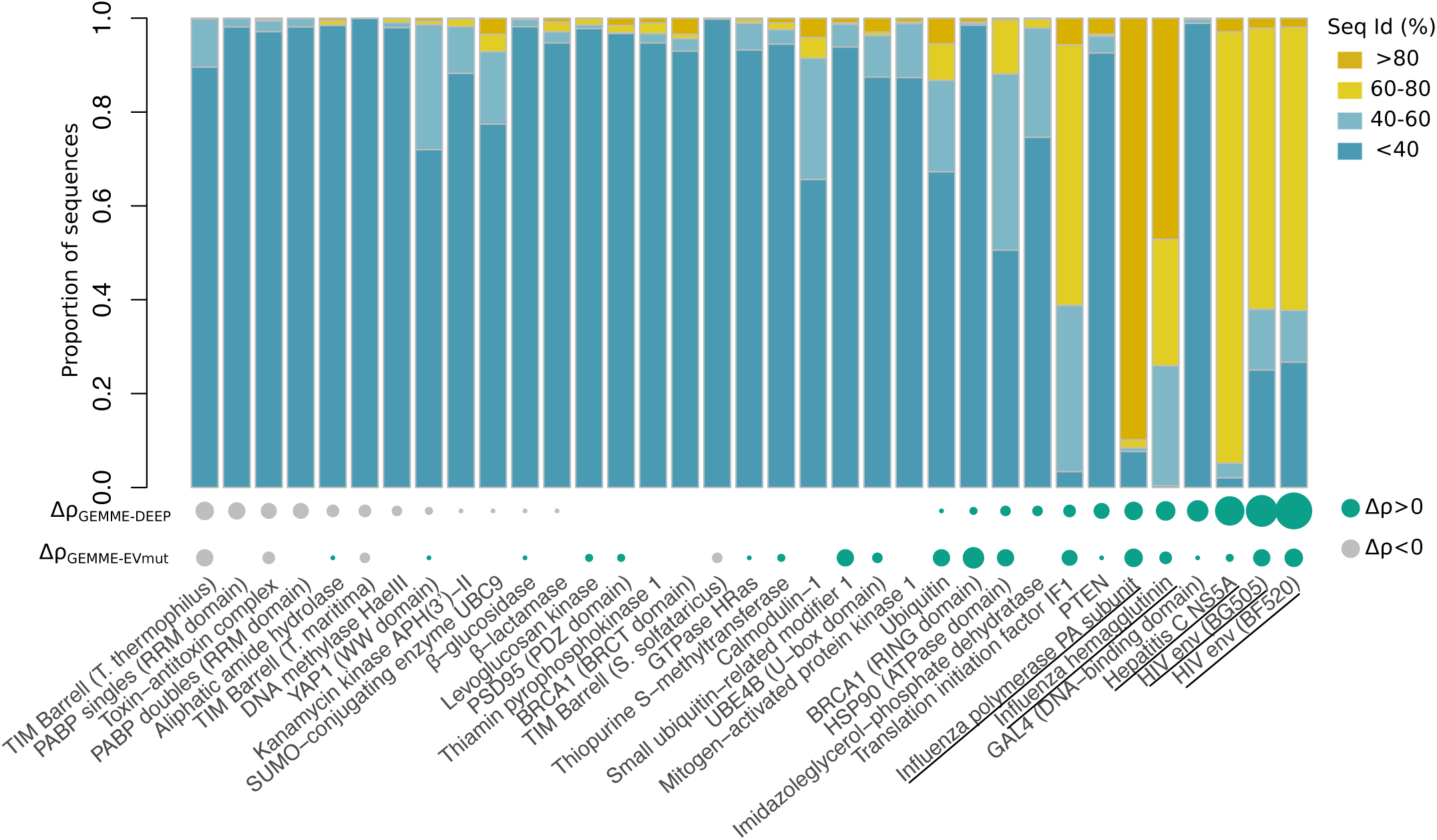
Sequence identities of the input alignments. The percentages of sequences sharing less than 40%, between 40 and 60%, between 60 and 80% and more than 80% with the query sequence are reported. Viral sequences are underlined. The dots at the bottom indicate the gain of performance of GEMME with respect to DeepSequence (Δ*ρ*_*GEMME–DEEP*_) and EVmutation (Δ*ρ*_*GEMMEEV-mut*_). The size is proportional to the absolute value of Δ*ρ* and the color depends on the sign. Δ*ρ* is positively correlated with the proportion of sequences sharing more than 60% identity with the query (Pearson correlation coefficient *R* = 0.71 for Δ*ρ*_*GEMME-DEEP*_ and *R* = 0.48 for Δ*ρ*_*GEMME-EV* *mut*_).

### Epistasis helps discriminate between rather frequent mutations

To compare different mutations occurring at the same position, GEMME’s model combines two contributions, namely the minimal evolutionary fit required to accommodate each mutation and the relative frequency of occurrence of the mutation (see *Methods*). The first term accounts for inter-site dependencies, and can thus be qualified as ‘epistatic’, whereas the second term is computed in a site-independent manner. If a mutation is rare and appears far away in the evolutionary tree, then both terms will be high and the mutation will be predicted as deleterious. On the contrary, if a mutation is frequent and found in sequences very close to the query, both terms will be small and hence, the predicted impact of the mutation will be small. The two terms will disagree in case of a rare mutation appearing in a sequence very similar to the query, or in case of a frequent mutation appearing only in highly divergent sequences. By default, GEMME puts a higher weight on the epistatic term (see *Methods*).

We systematically assessed the predictive power of each contribution taken separately (see *Methods*, **Fig. 4a** and **Supplementary Table S1**). Overall, the predictions issued by the epistatic contribution were found in better agreement with the experimental measures than those from the independent contribution (average Δ*ρ* = 0.04±0.09, median Δ*ρ* = 0.02). Yet, in about one third of the cases (11/35), it is better to rely on site-independent frequencies rather than evolutionary distances (**Fig. 4a**, on the right). Hence, the predictive power of the information coming from epistasis varies substantially from one protein to another. Such variability may arise from some intrinsic properties of the studied proteins, the experimental set up (*e.g.* measured phenotypes) and/or the properties of the input alignment. The bacterial DNA methyltransferase HaeIII and the nonstructural protein 5A (NS5A) from Hepatitis C virus provide two archetypal examples that help understand the influence of the input alignment. Both proteins display very contrasted performances between the two contributions, but while the epistatic term is the best one for the bacterial methyltransferase (**Fig. 4a**, on the left), the most accurate predictions for the viral NS5A are issued from the independent term (**Fig. 4a**, on the right). In the first case, a lot of mutations are rather frequent in the input alignment (**Fig. 4b**, on the left, in blue). More precisely, half of the mutations are at least 22% as frequent as the wild-type amino acid. By contrast, in the second case, half of the mutations are very rare (less than 0.03% as frequent as the wild-type amino acid) or simply not found in the alignment (**Fig. 4b**, on the left, in red). More generally, as the mutation average frequency of occurrence increases, so does the gain of the epistatic contribution over the independent one (**Fig. 4b**, in the middle, Pearson correlation coefficient *R* = 0.61). The shape of the mutation frequency distribution is also a good indicator (**Fig. 4b**, on the right, *R* = *-*0.60 with the distribution’s skewness). The best case scenario for the epistatic contribution is when a large body of mutations are rather frequent, and few mutations are rarer than average and widely spread (see the long left tail of the distribution in blue on **Fig. 4b**, on the left). Hence, when dealing with rather frequent mutations, looking at the overall divergence of the sequences where they appear, as quantified by our evolutionary fit, helps improving the discrimination between them.

**Figure 4:**
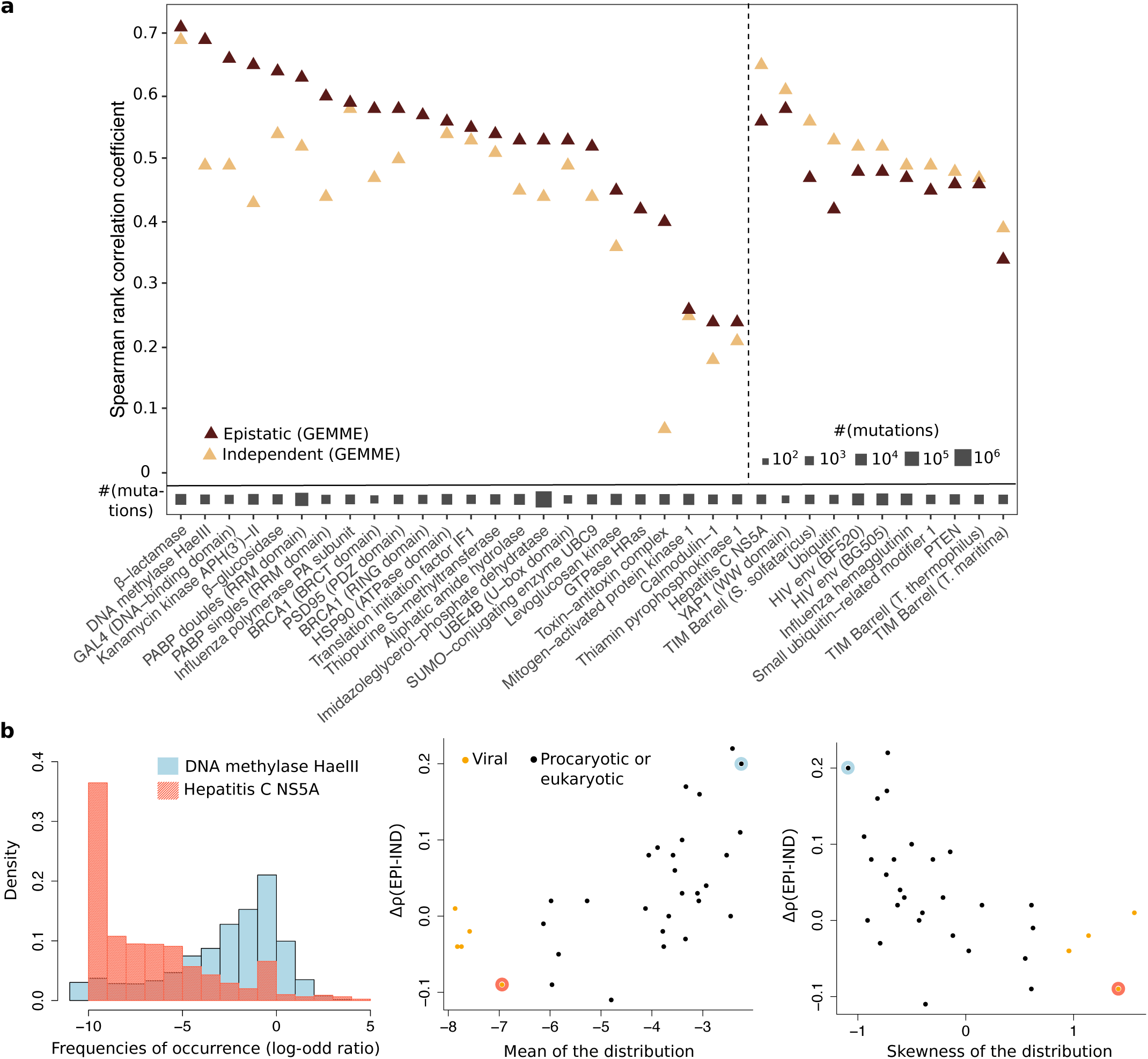
Comparison of predictive performances between the epistatic and independent contributions of GEMME’s model. **(a)** On the x-axis, proteins are divided in two groups according to the contribution yielding the highest correlation with experimental data (epistatic contribution on the left, independent one on the right) **(b) Left panel.** Examples of distributions for the mutations’ site-independent relative frequencies of occurrence. For each mutation, the reported value is the log-odd ratio between the number of sequences displaying the mutation over the number of sequences displaying the wild-type amino acid (see *Methods*). **Middle and right panels.** Difference in Spearman *ρ* coefficient in function of the mean (in the middle) and the skewness (on the right) of the log-odd ratio distribution. The skewness reflects the asymmetry of the distribution (positive skewness indicates a left tail while negative skewness indicates a right tail. The dots corresponding to the proteins taken as examples on the left panel are encircled.

### Evolutionary conservation is a valid proxy for position-specific sensitivity to mutations

Besides providing full protein mutational landscapes, mutational scans can be used to determine which positions in the protein are particularly sensitive to mutations. Such positions typically represent a small portion of the protein and can be viewed as its “weak” spots. The sensitivity of a position can be estimated by averaging its mutational outcomes over the 19 possible substitutions. In GEMME’s predictions, this average is strongly correlated to the position’s degree of conservation (**Supplementary Fig. S1**). This is expected as GEMME re-weights positions according to their evolutionary conservation to compare mutations occurring at different positions (see *Methods*). This results in highly conserved positions displaying overall higher predicted mutational effects than lowly conserved positions. We found that the conservation degrees alone provide estimates of position-specific sensitivities to mutations that are only slightly less accurate than the averages computed from GEMME’s full predicted matrices (**Fig. 5a**). This indicates that our conservation measure is already a good indicator of the extent to which a position will be sensitive to mutations. Importantly, this holds true even when the variability of the available sequence data is low. This means that we are able to capture meaningful signals in contexts where the content of information is poor. Moreover, our conservation measure compares well with the predictions issued by DeepSequence and EVmutation (**Fig. 5a**). In several cases, it better reflects experimentally determined mutational sensitivities than the averages computed from these predictions (**Fig. 5a**, points highlighted in blue).

**Figure 5:**
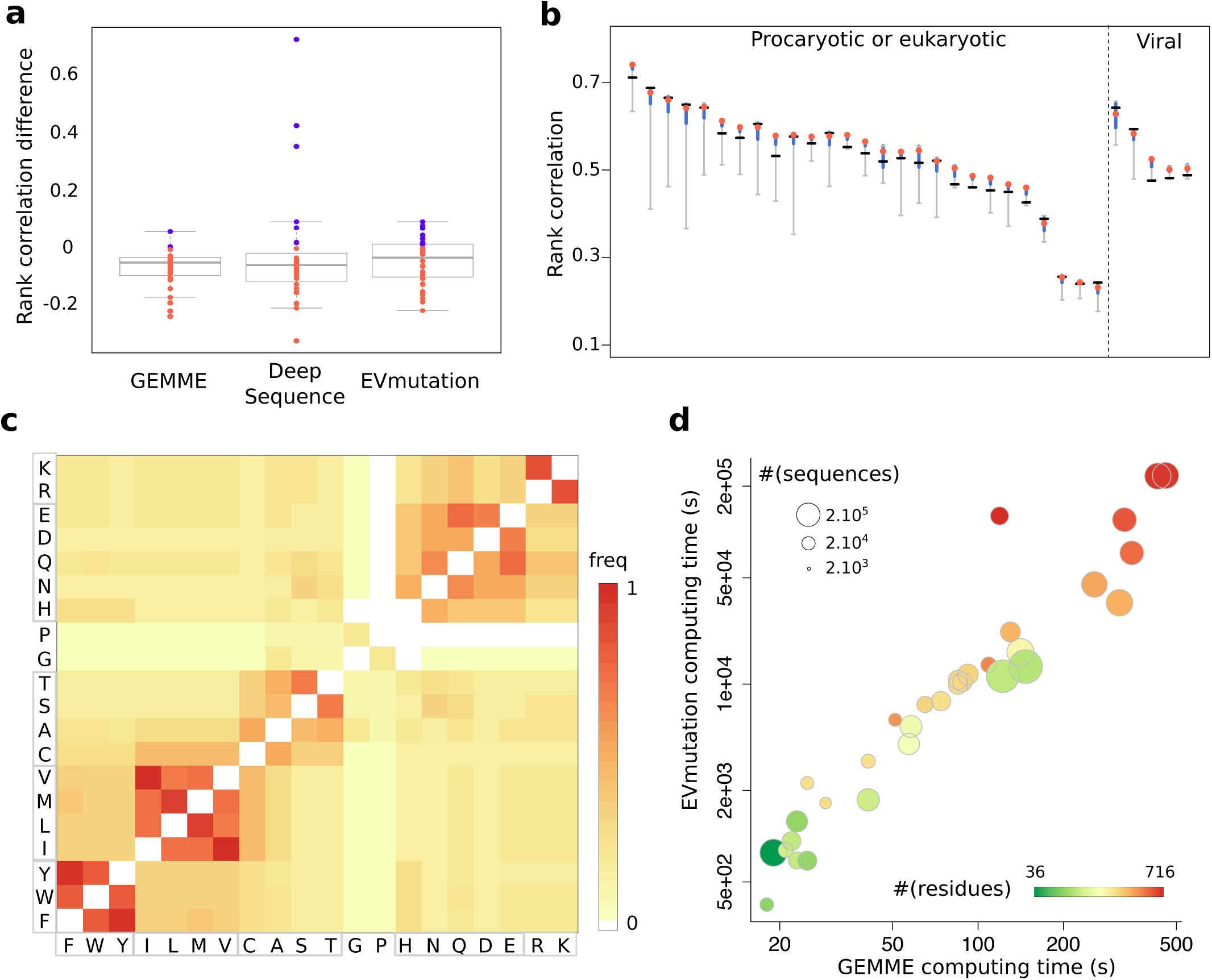
Analysis of GEMME’s parameters and computing time. **(a)** Differences in position rank correlation between the evolutionary conservation degrees computed by GEMME and the mutational effects predicted by GEMME, DeepSequence and EVmutation. Each points stands for a scan. Positive and negative differences are highlighted in blue and red, respectively. **(b)** Ranges of rank correlation obtained when varying the relative importance of GEMME’s independent and epistatic contributions. Each vertical segment corresponds to a deep mutational scan (same order as in **Fig. 2**). The red dot indicates the correlation obtained with GEMME’s default model, where the epistatic term is assigned a weight of 0.6 (and 0.4 for the independent term, see *Methods*). The black dash indicates the best performance achieved by the independent or epistatic contribution alone. The blue thick segments highlight the range of values obtained when varying the epistatic term’s weight between 0.5 and 0.8 (see also **Supplementary Fig. S2** and **S3**). **(c)** Amino acid grouping preferences observed in GEMME’s best performing models (parameters optimized for each scan). The color code goes from white (grouping never observed) to red (grouping observed for all scans). The amino acids are ordered so as to highlight the reduced alphabet used by default in GEMME. **(d)** Computing times of EVmutation and GEMME (in seconds, with logarithmic scales).

### GEMME’s results are robust and its model is transferable to other proteins

To assess the generalizability of GEMME’s model, we systematically evaluated the influence of its two main parameters on the quality of the predictions. The first parameter is the relative importance given to the epistatic and independent contributions. We observed that our default model, where the epistatic contribution is given more weight, systematically achieves similar or better correlation with experiments than each contribution taken alone (**Fig. 5b**, compare red dots and black dashes). In cases where the independent contribution performs better than the epistatic one, combining the two leads to only slightly lower performances. Moreover, varying the relative weights of the two terms around their default values has a very small impact on the quality of the predictions in most of the cases (**Fig. 5b**, blue segments). The second degree of freedom is the choice of the amino acid alphabet. GEMME relies on similarities between amino acids rather than identities to compute the mutations’ frequencies of occurrence. By default, we consider 7 classes of amino acids, grouping together the aromatic ones (F, W, Y), the hydroxyl-containing ones with alanine (C, A, S, T), the aliphatic hydrophobic ones (I, L, M, V), the positively charged ones (K, R) and the polar and negatively charged ones (H, N, Q, D, E). Glycine and proline are in a separate class each. We tested 164 different alphabet reduction schemes (see *Methods* and **Supplementary Table S3**) and found that the impact of the alphabet’s choice on the predictive performance is limited (average correlation standard deviation of 0.01, **Supplementary Fig. S2** and **S3**). So is the correlation gain obtained by optimizing the parameters for each scan (average Δ*ρ* = 0.02 ± 0.01, and median Δ*ρ* = 0.02, **Supplementary Fig. S4a**). Moreover, the amino acid grouping preferences exhibited by the best performing models are in good agreement with the alphabet chosen by default (**Fig. 5c**). This analysis shows that our results are robust to parameter changes and that our choices lead to predictions whose quality is close to the best one can hope for within GEMME’s framework. This is true overall and on most of the scans studied here, which makes us confident that our default model is directly transferable to other proteins.

### GEMME is faster than DeepSequence and EVmutation by several orders of magnitudes

To be applicable at large scale, computational scans should be fast. Given the input sequence data, it takes less than 10 minutes, on a single-core processor, for GEMME to generate any of the complete single-site mutational landscapes considered here. The corresponding proteins are of various lengths, comprising between 36 and 716 residues, and are associated with up to several hundreds of thousands of homologous sequences (**Fig. 5d**). By comparison, EVmutation requires several days of computation to deal with the biggest proteins of the dataset. Overall, GEMME is faster than EVmutation by a factor ranging between 19 and 1 072 (**Supplementary Table S2**). It should be stressed that EVmutation disregards some positions and some sequences from the input alignment while GEMME does not. DeepSequence is expected to be even more computationally expensive. Training one deep latent model on the *β*-lactamase family and computing the predictions required almost 7 hours (24 175 seconds) on a powerful graphics card (see *Methods*). This computing time has to be multiplied by 5 to obtain results similar to those reported in [3]. GEMME took only about 1 minute to treat the same protein on a single-core processor (**Fig. 5d** and **Supplementary Table S2**). Hence, we estimate DeepSequence to be several thousands times more computationally expensive than GEMME.

## Discussion

We have presented GEMME, a computational method for performing mutational scans of protein sequences. It exclusively exploits protein sequence data available in public databases. It relies on a few biologically sound assumptions about the relationship between protein sequence and function. Its algorithm is straightforward and requires setting only two parameters. It uses the mathematical tree structure underlying the evolution of natural sequences and it explores it by using simple new concepts (smallest path between wild-type and mutated sequences). This is markedly different from what has been developed previously. State-of-the-art methods feature many more parameters, infer some of them using sophisticated machine learning techniques, others empirically, pre-treat the input sequences to correct for bias, and do not explicitly model the evolutionary history relating these sequences. They are based on statistical inference and thus make assumptions on the space of all possible sequences. By contrast, GEMME directly exploits the information encoded in natural sequences. Despite its apparent simplicity, our method achieves similar or better performance than the state of the art, and it is the only viable option when dealing with viral sequences. It has the advantage of being faster than recently published methods by several orders of magnitude.

An important ingredient of GEMME is the inclusion of dependencies between the different positions in the sequence of interest. We are not the first ones to propose to account for ‘epistasis’ in the prediction of mutational effects. What has been done before was to explicitly model couplings between pairs of positions, inferred from co-occurring patterns in the input sequence data, or to implicitly model higher order dependencies by coupling each position to a ‘hidden’ variable. Let us stress that the introduction of such hidden variables do no ease interpretability. We adopt an orthogonal approach by introducing the notion of an evolutionary history relating the sequences observed today in nature. We infer such evolutionary history by quantifying global similarities between sequences, thus accounting for all positions and their inter-dependencies. Then we use the reconstructed evolutionary trees to identify functionally important positions, the rationale being that such positions should display a few amino acids that appeared and were fixed early in evolution.

We have implemented our method as a fully automated package and webserver, available at: www.lcqb.upmc.fr/GEMME/. It can deal with single mutations as well as combinations of mutations. It has been carefully evaluated against experimental measurements from 41 high-throughput mutational scans comprising between 313 and 496 137 data points. Moreover, we have provided an understanding of the contribution of ‘epistasis’ to the discrimination of mutations, based on the analysis of the variability of the input sequence data, and have demonstrated that our evolutionary-informed measure of conservation is a good indicator of the extent to which a position is sensitive to mutations. Finally, we have systematically assessed the influence of changing GEMME’s two parameters of the quality of the predictions. Our results are consistent and suggest that GEMME’s model is generalizable and transferable to other systems than the few tens studied here.

This work contributes to a better understanding and characterization of protein mutational landscapes. Such characterization is instrumental for the control of protein functions in physio-pathological contexts. Perspectives include the systematic description of the effects of mutations on protein interaction networks, a field that needs to expand in the coming years.

## Methods

### GEMME’s workflow

The GEMME (Global Epistatic Model for predicting Mutational Effects) method takes as input a multiple sequence alignment (MSA) in FASTA format, with the query sequence appearing on top. First, evolutionary conservation levels are computed using JET [22]. Then, GEMME predicts the mutational landscape of the query sequence. By default, it estimates the mutational outcomes of all possible single mutations. Alternatively, the user can provide an ensemble of single or multiple mutations of interest.

#### Homologous sequences retrieval and selection

The user can ask GEMME to compute conservation levels directly on the input MSA. Alternatively, GEMME will automatically launch a PSI-BLAST [27] search to retrieve up to 5 000 sequences related to the query. Then, a number of selection criteria will be applied to filter the set of related sequences. By default, sequences redundant with the query (*>*98% identity) or too far (*<*20% identity), too small (*<*80% coverage), too gapped (*>*10% of the size of the alignment) or not significant enough (e-value ≥10^*-*5^) are removed. If the number of remaining sequences is too low (*<*100), the selection criteria are progressively relaxed as described in [22]. All parameters are adjustable by the user.

#### Evolutionary conservation levels (T_*JET*_ )

The calculation of evolutionary conservation levels relies on a Gibbs-like sampling of the filtered set of related sequences [22]. Sequences are classified into four groups, depending on the degree of identity they share with the query (20-39%, 40-59%, 60-79%, and 80-98%). *N* sequences are randomly picked up from the four classes to construct a subset representing the diversity of the whole set. The sequences are then aligned and a distance tree is constructed from the alignment using the Neighbor-Joining algorithm [28]. For each position in the query sequence, a *tree trace level* is computed: it corresponds to the level *l* in the tree where the amino acid at this position appeared and remained conserved thereafter (see [22] for a more precise definition). Let us recall that this definition of evolutionary trace is notably different from the measure defined by Lichtarge and co-authors to rank protein residues [20, 21].

This procedure is repeated *N* times and the *tree trace levels* are averaged over the *N* trees to get more statistically significant values, which we denote *relative trace significances*, or *T*_*JET*_, and which are expressed as [22]:

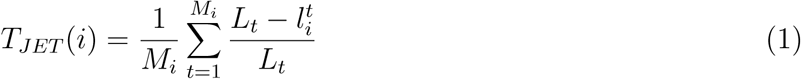

where 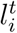 is the *tree trace level* of residue *r*_*i*_ in tree *t, L*_*t*_ is the maximum level of *t* and *M*_*i*_ is the number of trees where a non null *tree trace level* was computed for *r*_*i*_. *T*_*JET*_ values vary in the interval [0,1] and represent averages over all trees of residues’ evolutionary conservation levels.

To produce the results reported here, we used the most recent version of the JET method, namely JET^2^ [29] (available at: www.lcqb.upmc.fr/JET2). JET^2^ uses MUSCLE [30] to align sequences. JET^2^ was launched in its iterative mode: the procedure described above was repeated 10 times and the maximum conservation value obtained over the 10 runs was retained for each residue.

#### Predicted effects: comparison of mutations occurring at the same position

To compare mutations occurring at the same position, GEMME combines two contributions. The first one is termed *epistatic* and corresponds to the minimal evolutionary fit required to accommodate the mutation of interest. The second one is termed *independent* and reflects the relative frequency of occurrence of the mutation. Hence, the predicted effect of a mutation X-to-Y at position *i* is expressed as:

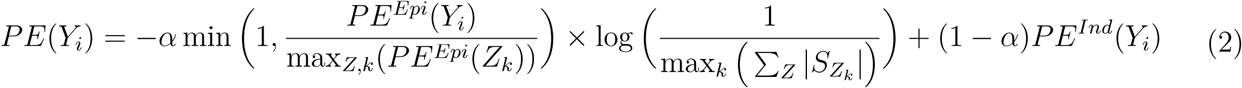

where *PE*^*Epi*^(*Y*_*i*_) and *PE*^*Ind*^(*Y*_*i*_) are the values of the epistatic and independent contributions (defined below), respectively. The term min 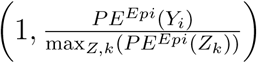 scales the value of the epistatic contribution between 0 and 1. The maximum value max_*Z,k*_(*PE*^*Epi*^(*Z*_*k*_)) is determined over all positions *k* and all possible substituting amino acids *Z* for a given protein. The term 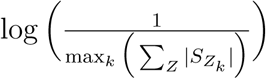, where 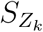 is the ensemble of sequences displaying Z at position *k*, gives the lowest possible log-ratio value. It corresponds to the extreme case where all non-gapped sequences at position *k* display the wild-type amino acid. It is used as a multiplying factor here so as to be able to combine the predictions coming from the two different contributions. The coefficient *α* determines the relative weight of each contribution. By default, *α* is set to 0.6, and sequence counts are calculated using a reduced representation of the amino acid alphabet comprised of seven classes: FWY, ILMV, CAST, G, P, HNQDE, RK (LZ-BL-7 in **Supplementary Table S3**).

#### Epistatic contribution (*PE*^*Epi*^)

The evolutionary relationships between all the sequences in the input MSA can be represented by a tree, which is not explicitly computed here. The topology of that tree is implicitly reflected by the *T*_*JET*_ values, which were computed and averaged over many small trees. We illustrate this by considering 2 positions *i* and *j* at which the query sequence *q* displays S and G, respectively (**Fig. 1a-b**). Position *i* is lowly conserved (*T*_*JET*_ =0.2) while position *j* is highly conserved (*T*_*JET*_ =0.8). This implies that *q* belongs to a smaller subtree of sequences displaying S at position *i* (**Fig. 1b**, on the left, dark grey rectangle), and to a bigger subtree of sequences displaying G at position *j* (**Fig. 1b**, on the right, light grey rectangle).

To estimate how close some sequence *s* is from the query sequence *q*, we define the evolutionary distance *D*_*evol*_(*q, s*) as:

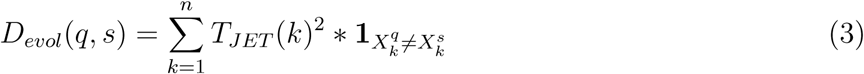

where *n* is the length of *q*, 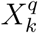 is the amino acid of *q* at position *k*, and 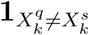 is the indicator function. Only positions where the amino acid in *s* is different from the amino acid in *q* (**Fig. 1c**, in red) contribute to the sum, and the level of contribution depends on the level of conservation of the position (**Fig. 1c**, second color strip).

To assess the effect of a mutation X-to-Y at position *i* in *q*, we select the subset 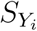 of sequences displaying the mutation, and look for the sequence within 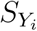 being the closest to *q*. The resulting minimal evolutionary distance estimates how far from *q* one has to go in the tree to observe a sequence bearing Y at *i*. Hence, the predicted effect of mutation Y at position *i, PE*^*Epi*^(*Y*_*i*_), is expressed as:

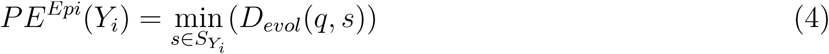

To avoid bias due to the presence of a peculiar sequence or of an alignment error in the MSA, we require that there exists at least one sequence different from the closest one and at a similar distance to the query. For this, we rank all evolutionary distances in ascending order and compute the difference between the first and second ones. If the difference is lower than an arbitrarily chosen cutoff of 5, then we keep the first one. Otherwise, we replace it by the second one. In cases where the ensemble *S*_*Y*_ is empty (*i.e.* no sequence displaying Y at position *i* could be found), then we set *PE*^*Epi*^(*Y*_*i*_) = +∞.

This metric enables to directly compare and rank several substitutions at a given position. Given two amino acids Y and Z substituting X at position *i*, one can express the difference *PE*^*Epi*^(*Y*_*i*_) *-PE*^*Epi*^(*Z*_*i*_) as:

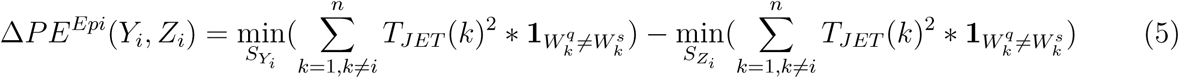

where the sums are computed over all positions except *i*, as the contribution of *i* cancels out. Consequently, this difference quantifies how much the global sequence context of position *i* has to change to accommodate the X-to-Y substitution versus X-to-Z. The substitution displaying the highest change will be predicted as the most deleterious one at that position.

As an example, let us consider mutations S-to-T and S-to-A at position *i* on **Figure 1a-b**. The S-to-T mutation induces a smaller minimal amount of changes than the S-to-A mutation, as the closest sequence displaying T at *i* (indicated by a blue star on **Fig. 1b**, left panel) has a lower evolutionary distance to the query than the closest sequence displaying A at *i* (green star). The S-to-A substitution at *i* systematically implies a G-to-V mutation at another position *j*, while the S-to-T substitution does not.

#### Independent contribution (*PE*^*Ind*^)

This contribution focuses only on the position where the mutation occurs. Hence, the effect of a substitution X-to-Y at positions *i* will be estimated as:

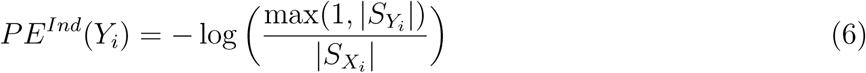

where 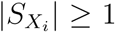 since at least the query sequence *q* displays X at position *i*, and the value 1 in the numerator serves as a pseudo-count in case 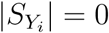, *i.e.* no observed sequence displays Y at position *i*. According to this model, the fewer sequences displaying the mutation, the more deleterious the mutation, for a given position.

#### Normalized predicted effects: comparison of mutations occurring at different positions

To compare substitutions at different positions in the query protein, we proceed through a normalization step. The normalized predicted effect (NPE) of a mutation X-to-Y at position *i* is expressed as:

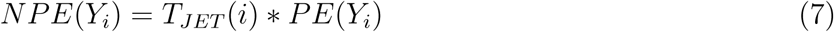

where *PE*(*Y*_*i*_) can be epistatic or independent. The normalization will result in highly conserved positions being highly intolerant to mutations while any substitution at a poorly conserved position will have a small effect (compare the two matrices on **Figure 1c**, where the highly conserved positions are highlighted by arrows).

In the toy example pictured on **Figure 1a-b**, the evolutionary distance to the closest sequence displaying A at position *i* (indicated by a green star on **Fig. 1b**, on the left) is slightly longer than that to the closest sequence displaying V at position *j* (**Fig. 1b**, on the right, orange star). Nevertheless, the normalization step will result in mutation G-to-V at *j* being predicted as more deleterious than mutation S-to-A at *i* since position *j* is far more conserved than position *i*.

#### Extended global epistatic model for multiple mutations

We extended the global epistatic model to deal with combinations (pairs, triplets…) of mutations. The normalized predicted effect of a given combination of *p* mutations is expressed as:

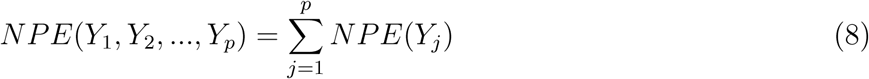

#### Parameters set up

To determine the default value of *α* and the default reduced amino acid alphabet scheme, we systematically computed predictions for all values of *α*, ranging between 0 and 1 by increments of 0.1, and for 164 reduced alphabets (see below and **Supplementary Table S3**). For each combination (*α*,alphabet), we computed its mean squared displacement from the best performing combination. Among the 5 combinations displaying the lowest mean squared displacements, we chose the combination with the lowest median squared displacement, namely *α* = 0.6 and LZ-BL-7 as the alphabet scheme. To identify the model yielding the best performance, for each experimental scan, the coefficient *α* was varied between 0 and 1 by increments of 0.1, and the 164 amino acid alphabets were systematically tested.

### Experimental datasets and input alignments

To assess GEMME’s performance and compare it fairly with DeepSequence and EVmutation, we considered the same body of experimental data and the same input alignments as those reported in [3] (see [31, 32, 33, 34, 35, 4, 36, 37, 38, 39, 40, 41, 42, 43, 44, 45, 46, 47, 48] for details about each experiment). We computed GEMME’s predictions for 41 deep mutational scans across 34 full proteins, protein domains or protein complexes. Among those, 38 scans comprise only single-site mutations and 3 scans comprise combinations of mutations. Two scans are associated to 2 different domains of the same protein (BRCA1), and one scan is associated to a protein complex (toxin-antitoxin complex). In the main text and figures, we report the performance obtained against one measured phenotype from one scan, for each protein. In case of multiple scans associated to the same protein, we focus on the most recent one. There is one exception, namely PABP, for which 2 scans were retained because one deals with single-site mutations and the other one with multiple mutations. The selected measured phenotype is the one yielding the best agreement with the predictions. All results for all measured phenotypes from all scans are reported in **Supplementary Table S1**.

### Reduced amino acid alphabets

A reduced alphabet is a clustering of amino acids based on their relative similarity. We tested 164 different alphabet schemes [49], whose names and characteristics are reported in Supplementary Table S3. They comprise between 2 and 19 letters. AB schemes were defined based on the ability of standard methods to correctly predict secondary structure from the simplified sequences [50]. CB schemes were produced by using the Miyazawa-Jernigan interaction matrix [51] and a distance-based clustering scheme [52]. DSSP and GBMR schemes were designed to maximally preserve structural information [53]. HSDM and SDM schemes were defined based on new substitution matrices derived from structural alignments of proteins with low-sequence identity [54]. LR is a 10-letter alphabet intended to increase the sensitivity of protein alignment searches [55]. LW-I and LW-NI schemes were designed to preserve information in global sequence alignments between a sequence and its reduced-alphabet version [56]. Notice that LW-I and LW-NI are identical at the level of 2, 3 and 15 through 19 letters, and that CB and LW are identical at the 2-letter level. LZ-MJ and LZ-BL were defined based on the identification of deviations of pair frequency counts from a random background, computed on the Miyazawa-Jernigan and BLOSUM 50 matrices [57]. ML schemes are based on the BLOSUM 50 substitution matrix [58]. MM is a 5-letter alphabet based on the Johnson-Overington matrix [59] which proved useful for aligning homologous sequences and assessing folds [60]. MS is a 6-letter alphabet based upon intuition and a study of the effects of disulfide bonds on protein folding which suggested separating aliphatic hydrophobic and aromatic hydrophobic residues [61]. TD schemes are based on intuitive physicochemical considerations [62]. WW is a 5-letter alphabet derived from the Miyazawa-Jernigan matrix by preserving maximal similarity between a reduced-alphabet version of the matrix and the full 20 *×* 20 matrix [63].

### Comparison of performance

Predictions for DeepSequence and EVmutation were directly taken from [3]. To evaluate and compare performances, we used Spearman rank correlation coefficient *ρ* as the primary metric. This choice was also made in previous studies [3, 7, 8] and is justified by the fact that we do not expect a linear relationship between predicted and experimental values. For each protein, correlations were computed on the set of mutations for which both experimental measures and predictions from DeepSequence and EVmutation were available.

### Comparison of computing times

To measure GEMME’s computing time, we ran the tool starting from the input MSA and the result of the PSI-BLAST search. Hence, for each protein, we measured the elapsed (wall clock) time required to compute the conservation levels and the predictions from all models (default, independent and epistatic). The computation of the conservation levels was realized by running one iteration of JET^2^. The resulting matrices of predicted mutational effects were very similar (99.6 ± 0.4 % Pearson correlation coefficient on average) to those obtained when running 10 iterations of JET^2^ and retaining the maximum conservation level over the 10 runs. EVmutation’s code was downloaded from https://github.com/debbiemarkslab/EVmutation and https://github.com/debbiemarkslab/plmc. For each protein, we measured the elapsed (wall clock) time required to compute the pairwise couplings and the predictions from both the independent and epistatic models. The same input MSAs, taken from [3], were given to GEMME and EVmutation. In these alignments, some positions are flagged because they are highly gapped. GEMME considered all positions whereas EVmutation disregarded the flagged ones. EVmutation also disregarded highly gapped sequences. The numbers of positions and sequences considered by each tool are reported in Table S2. Calculations were realized on a single core processor Intel Xeon E5-2630 v4 @ 2.20GHz. DeepSequence’s code was downloaded from https://github.com/debbiemarkslab/DeepSequence. We ran the tool to train one model on the input MSA for the *β*-lactamase and compute the predictions. The calculation was realized on a NVIDIA TITAN Xp graphics card. We tried and ran the tool on a single core processor but it produced an error.

## Acknowledgements

We thank Tristan Bitard-Feildel for providing computing time for DeepSequence, and Anne Lopes and Hugues Richard for careful reading of the manuscript.

## SUPPLEMENTARY MATERIAL

Table S1: **Detailed predictive performances. (see supTab allResults.xlsx)**

**Table S2:** Comparison of computation time between GEMME and EVmutation.

| Protein | GEMME |  |  | EVmutation |  |  |
| --- | --- | --- | --- | --- | --- | --- |
| | $\#(res)$ | $\#(seq)$ | Elapsed Time | $\#(res)$ | $\#(seq)$ | Elapsed Time |
| IF1 | 72 | 9 190 | 18s | 69 | 9 131 | 5m 55s |
| YAP1 | 36 | 86 353 | 19s | 30 | 86 271 | 12m 59s |
| POLG | 114 | 11 423 | 21s | 113 | 10 473 | 13m 30s |
| SUMO1 | 101 | 21 695 | 22s | 76 | 21 479 | 15m 28s |
| RL401 | 76 | 33 026 | 23s | 71 | 32 741 | 20m 51s |
| UBE4B | 104 | 16 478 | 23s | 75 | 16 411 | 11m 32s |
| GAL4 | 75 | 22 985 | 25s | 62 | 22 980 | 11m 31s |
| BRCA1 (BRCT) | 237 | 8 391 | 25s | 186 | 8 327 | 37m 17s |
| TPMT | 245 | 6 688 | 29s | 177 | 6 666 | 27m 32s |
| BRCA1 (RING) | 110 | 39 396 | 41s | 75 | 39 123 | 28m 55s |
| TPK1 | 243 | 9 966 | 41s | 201 | 9 923 | 51m 54s |
| PTEN | 403 | 8 566 | 51s | 304 | 8 474 | 97m 3s |
| UBC9 | 158 | 32 486 | 57s | 138 | 32 223 | 67m 17s |
| CALM1 | 149 | 36 224 | 58s | 139 | 35 850 | 87m 44s |
| BLAT | 263 | 14 783 | 1m 5s | 252 | 14 689 | 122m 24s |
| HSP82 | 240 | 23 447 | 1m 14s | 218 | 23 244 | 128m 50s |
| TIM (*S. solf*) | 248 | 23 743 | 1m 25s | 237 | 23 683 | 165m 24s |
| TIM (*T. mari*) | 252 | 23 745 | 1m 25s | 239 | 23 683 | 177m 39s |
| TIM (*T. ther*) | 254 | 23 742 | 1m 28s | 236 | 23 680 | 172m 10s |
| KKA2 | 264 | 29 808 | 1m 32s | 226 | 29 705 | 192m 55s |
| B3VI55 | 439 | 12 925 | 1m 49s | 364 | 12 880 | 223m 6s |
| PA | 716 | 19 611 | 1m 59s | 716 | 18 527 | 2 126m 11s |
| DLG4 | 101 | 210 888 | 2m 2s | 82 | 210 559 | 187m 45s |
| MTH3 | 330 | 26 513 | 2m 10s | 318 | 26 431 | 367m 12s |
| RASH | 189 | 84 762 | 2m 21s | 164 | 83 880 | 270m 25s |
| PABP | 96 | 246 405 | 2m 27s | 79 | 245 601 | 217m 11s |
| MK01 | 360 | 65 626 | 4m 17s | 288 | 64 888 | 753m 33s |
| AMIE | 346 | 76 372 | 5m 15s | 247 | 76 177 | 571m 5s |
| HG | 565 | 51 012 | 5m 28s | 544 | 47 013 | 2 005m 16s |
| BG | 501 | 49 471 | 5m 48s | 441 | 49 112 | 1 213m 58s |
| BG505 | 686 | 73 877 | 7m 10s | 653 | 71 227 | 3 881m 3s |
| BF520 | 686 | 73 441 | 7m 38s | 657 | 70 797 | 3 917m 12s |

GEMME considers the whole input alignment whereas EVmutation removes very gapped columns and sequences. This results in EVmutation treating a smaller number of residues ( $\#(res)$ ) and a smaller number of sequences ( $\#(seq)$ ) compared to GEMME (see *Methods*). The 32 proteins are ordered by increasing GEMME's computing time.

**Table S3:**
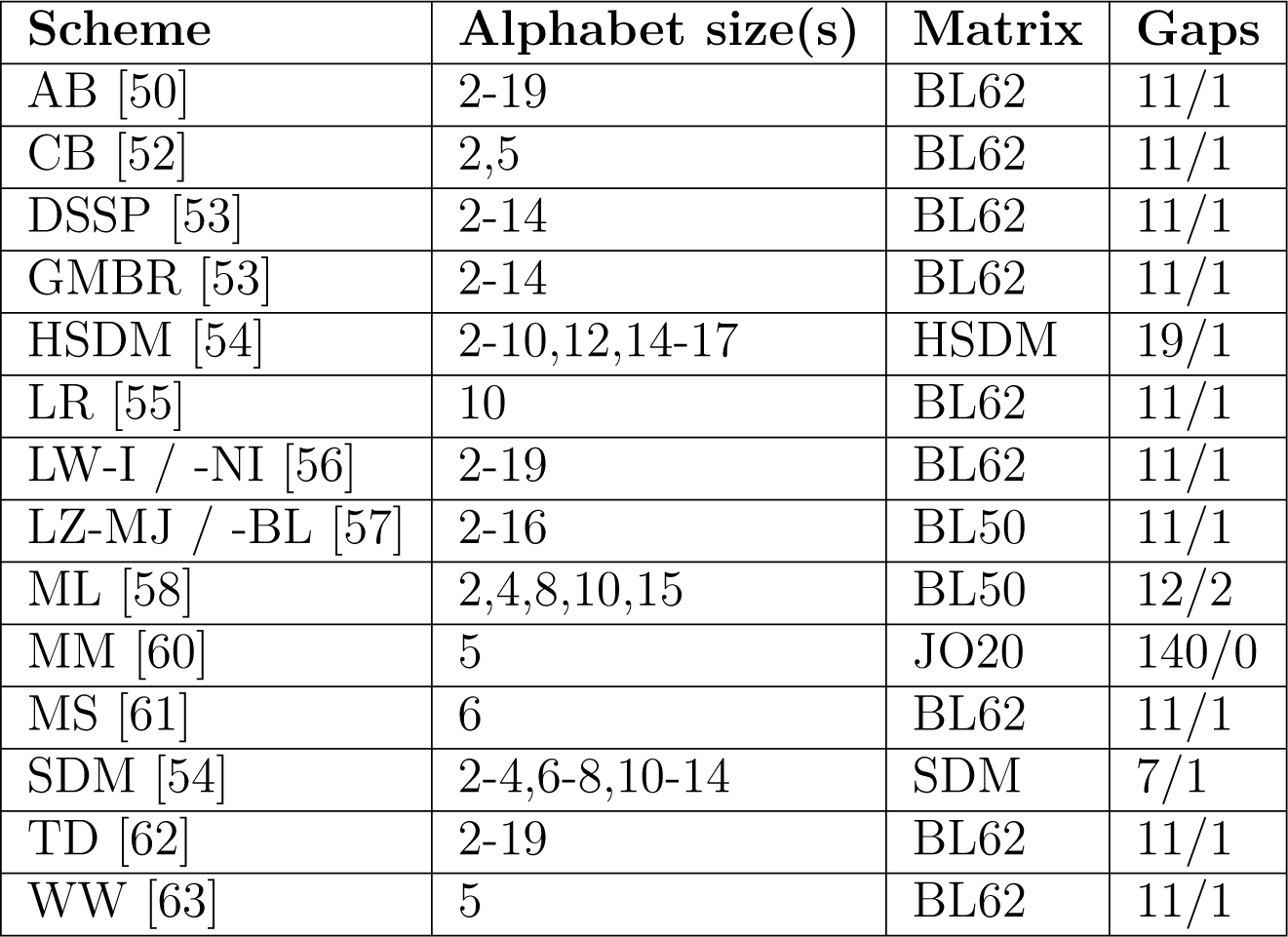
Characteristics of the investigated reduced alphabets. Abbreviations and references are listed in the first column. Alphabet sizes, matrix used and gap penalties are also given. BL50 and BL62: BLOSUM matrix [64] with 50 and 62% identity cutoff. JO20: Johnson-Overington matrix [59].

**Figure S1:**
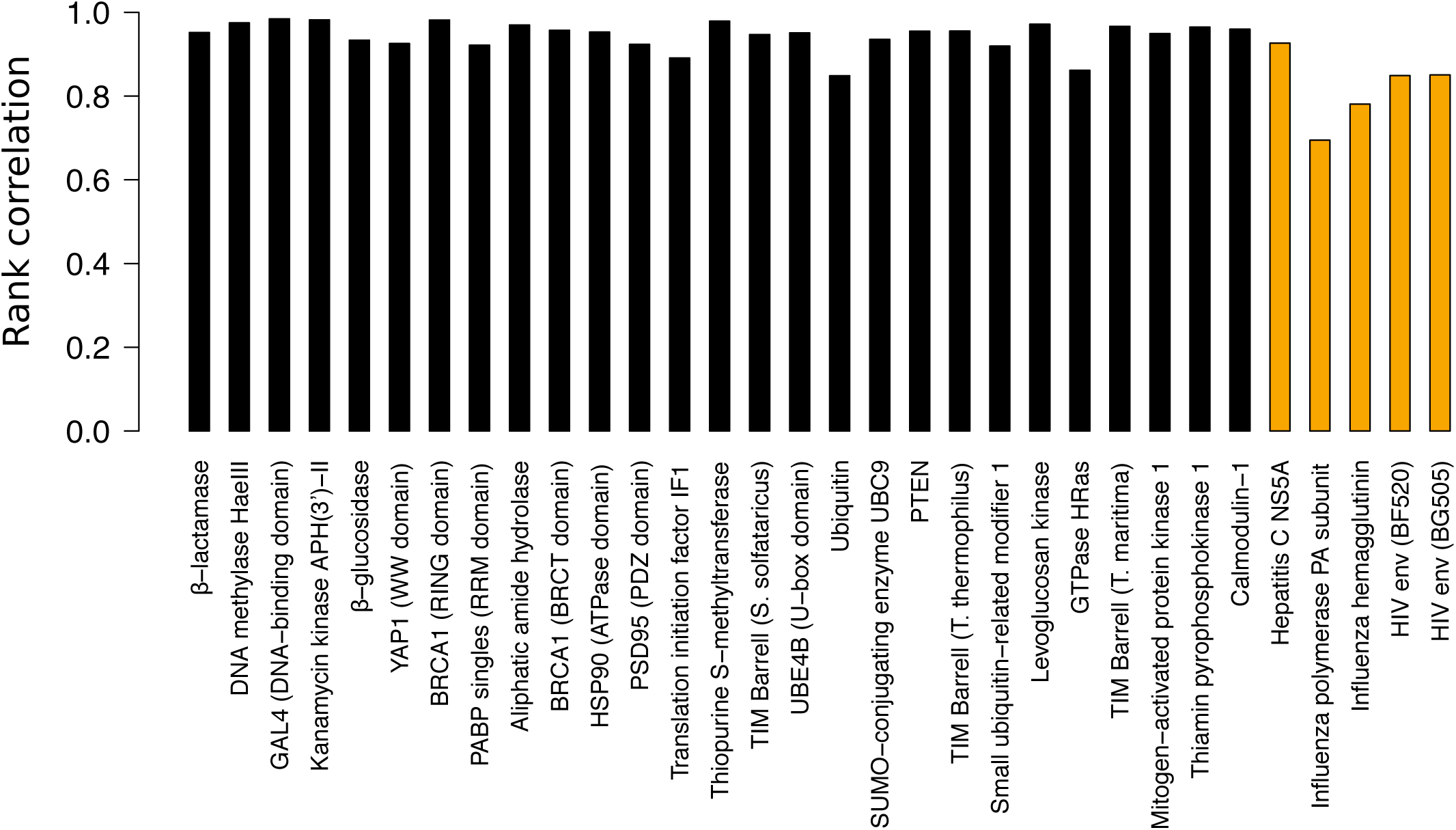
Rank correlations between GEMME’s predicted position-specific mutational sensitivities and evolutionary conservation levels. Viral proteins are highlighted in orange.

**Figure S2:**
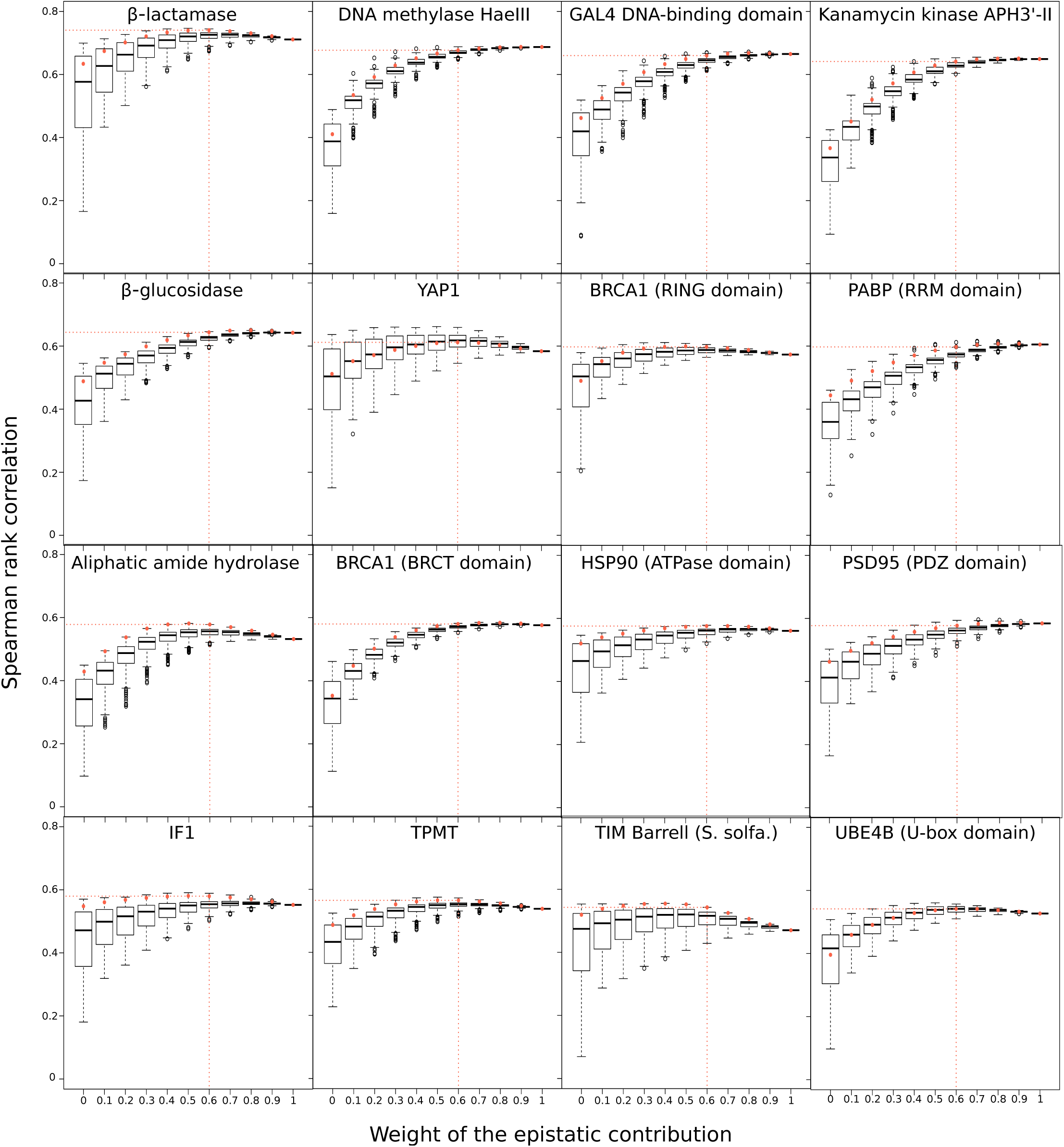
Distributions of Spearman rank coefficients. For each value of the epistatic contribution weight, the boxplot gives the distribution of *ρ* computed against experimental measurements for the 164 tested reduced amino acid alphabet. The red dot corresponds to the LZ-BL-7 alphabet, used by default in GEMME.

**Figure S3:**
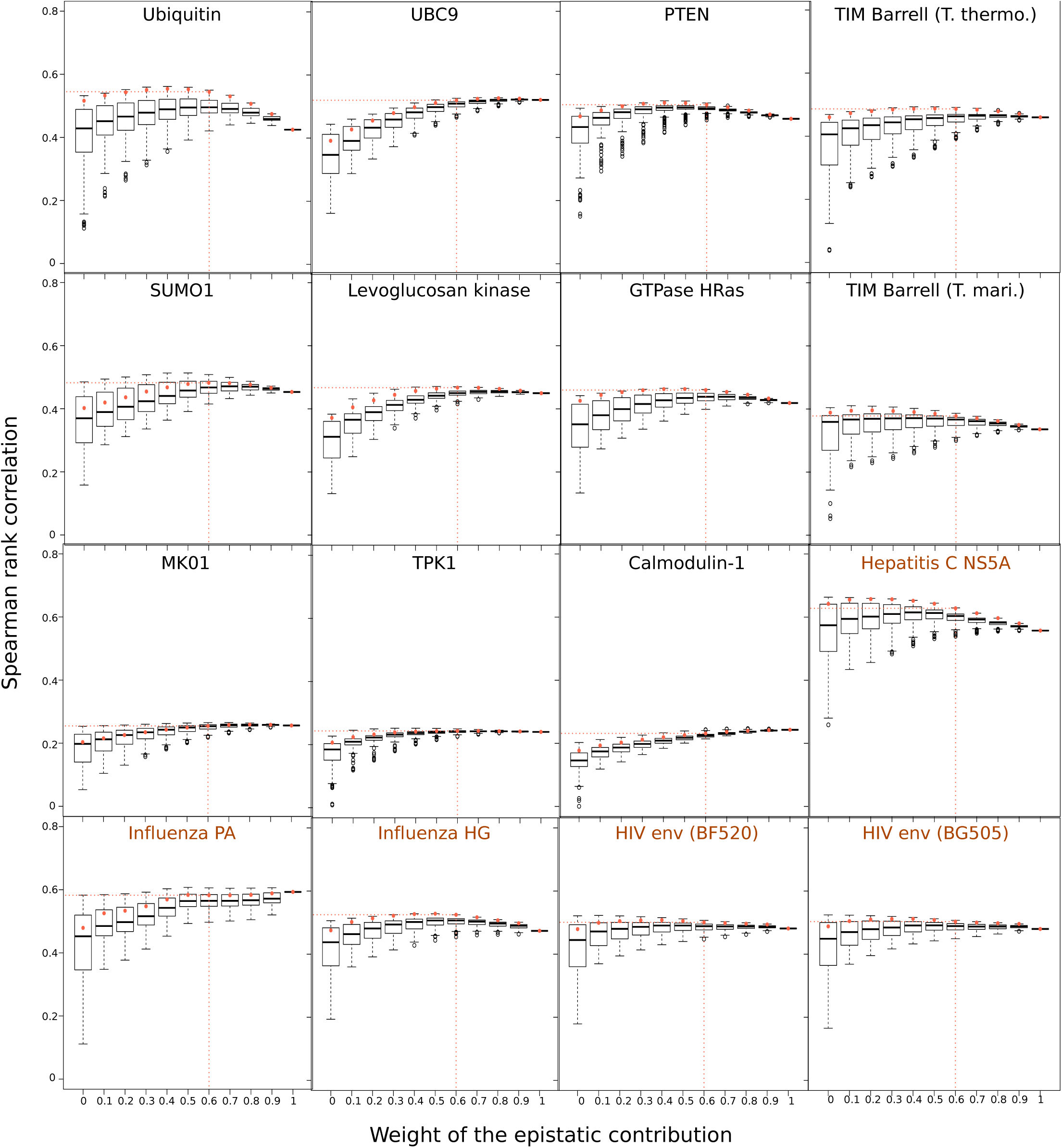
Distributions of Spearman rank coefficients. For each value of the epistatic contribution weight, the boxplot gives the distribution of *ρ* computed against experimental measurements for the 164 tested reduced amino acid alphabet. The red dot corresponds to the LZ-BL-7 alphabet, used by default in GEMME. Viral proteins are highlighted in orange.

**Figure S4:**
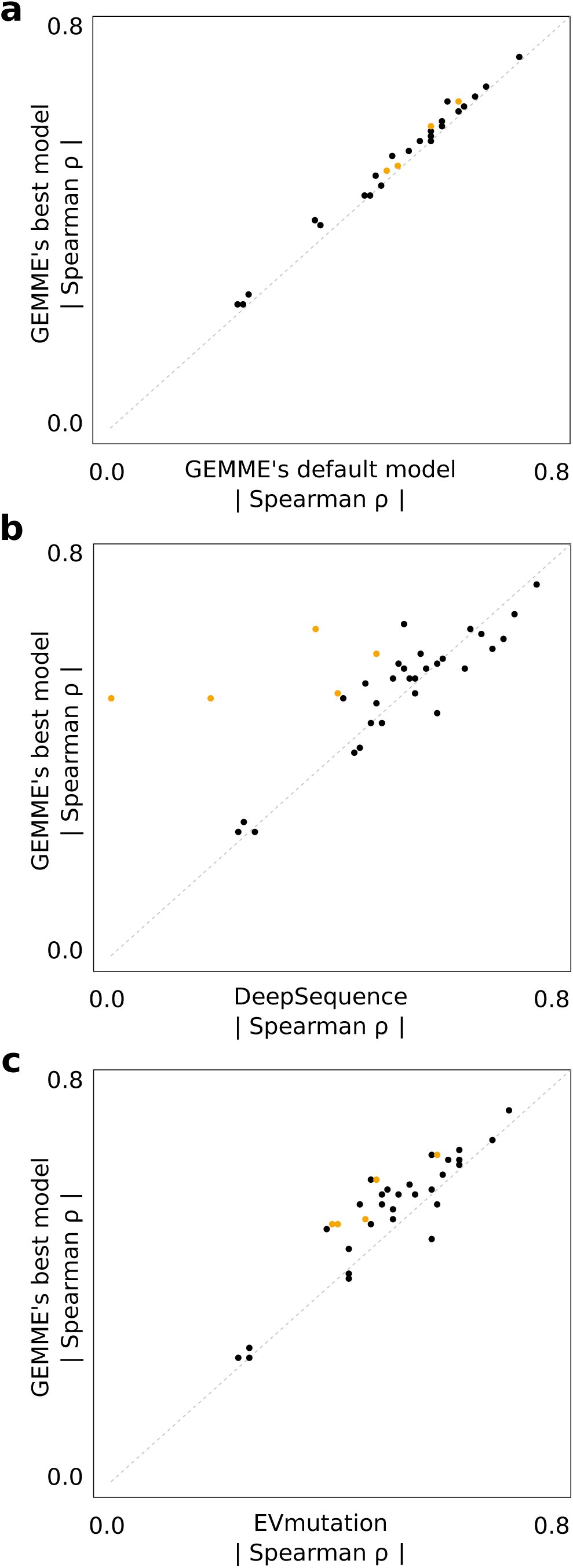
Comparison of predictive performances. GEMME’s best model, determined for each scan, is compared with GEMME’s default model (a), DeepSequence (b) and EVmutation’s epistatic model (c). Spearman rank correlation coefficients *ρ* were computed between predicted and experimental measures for 35 experiments corresponding to 34 proteins.

